# Multimerization of HIV-1 integrase hinges on conserved SH3-docking platforms

**DOI:** 10.1101/301721

**Authors:** Meytal Galilee, Akram Alian

## Abstract

New anti-AIDS treatments must be continually developed in order to overcome resistance mutations including those emerging in the newest therapeutic target, the viral integrase (IN). Multimerization of IN is functionally imperative and provides a forthcoming therapeutic target. Allosteric inhibitors of IN bind to non-catalytic sites and prevent correct multimerization not only restricting viral integration but also the assembly and maturation of viral particles. Here, we report an allosteric inhibitor peptide targeting an unexploited SH3-docking platform of retroviral IN. The crystal structure of the peptide in complex with the HIV-1 IN core domain reveals a steric interference that would inhibit conserved docking of SH3-containing domain with the core domain vital for IN multimerization, providing a template for the development of novel anti-IN allosteric inhibitors.

---

In the absence of a curative treatment, the highly active antiretroviral therapy (HAART) keeps the HIV-1 virus of AIDS patients under control. HAART combines drugs targeting different stages of viral replication including the integration step catalyzed by the integrase protein (IN) (1). Integration of viral DNA into host genome involves two steps catalyzed by IN: (i) cleavage of a dinucleotide from each 3’-end of the viral DNA (3’-processing), and (ii) insertion of this processed viral DNA into the host DNA (strand-transfer) (2). Clinical IN strand transfer inhibitors (INSTIs) target the catalytic site of the enzyme to specifically inhibit the DNA joining reaction, however, as with all anti-AIDS treatments, the continued success of these drugs is persistently disrupted by resistance mutations (1,2). Although 3′-processing can be carried out by monomeric IN (3), the assembly of IN functional multimers is imperative for the strand-transfer activity (4–8), and for virus particle maturation and production (reviewed in (9,10)). In the continued quest to identify and develop new drugs, allosteric inhibitors that bind sites outside the catalytic core and disrupt IN multimerization are emerging with potent therapeutic potential (11–14).

Tetramers of IN are formed by the reciprocal swapping of the three, N-terminal, catalytic core and C-terminal, canonical domains of IN. The two internal IN protomers, where catalytic binding of viral and host DNA takes place, make up the majority of the tetramer interface. The outer two protomers, and other surrounding units in higher order assemblies, provide supportive domains indispensible for the assembly of the tetrameric cores (4–8). We recently showed that targeting the interface between the N-terminal domain (NTD) and catalytic core domain (CCD) of IN, using a specific antigen-binding fragment (Fab), inhibits IN tetramer formation and consequent enzymatic activity and virus particle production (9). Disrupting the dimerization of the elementary dimeric blocks building IN tetramers has also been explored as a strategy to inhibit IN activity (15).

Hindering the assembly of IN functional multimers is only one side of the coin. Allosteric interference has also been shown to promote the formation of aberrant IN multimers and aggregates. The potential of allosteric IN inhibitors has been demonstrated through the thorough characterization of the “LEDGF pocket” formed at the dimer interface of IN and the development of LEDGIN (or ALLINI) inhibitors that bind to it (11) (Figure 1A). Although less investigated, other IN pockets capable of allosteric inhibitor binding have also been identified (Figure 1A): binding of the “Y3” molecule to a pocket near the N-terminal end of CCD α-helix 4, designated Y3-pocket, has been shown to inhibit 3’-processing and strand-transfer activities (16); the “sucrose” binding pocket found along the CCD dimer interface and flanked by two LEDGF pockets (17,18) has recently been targeted by the natural product kuwanon-L, which inhibited IN activity in a pattern similar to LEDGINs (19). Another fragment-binding pocket “FBP” has also been identified at the CCD dimer interface (20). Although inhibitory profiles for this FBP pocket have not been shown, our previous work on IN from the feline virus showed that a single Phe187Lys mutation (Phe185 in HIV-1 IN) around this site inhibits dimer formation (3). Fragment-based screening and structural studies have revealed the capability of all three pockets (Y3, FBP and LEDGF) to bind numerous small molecules (21). Most recently, a new class of LEDGINs has been shown to distinctively bind to undefined interfaces of both CCD and CTD of IN (22). Therefore, IN harbors several allosteric hotspots that can be explored for novel anti-multimerization intervention strategies.

**Figure 1.**
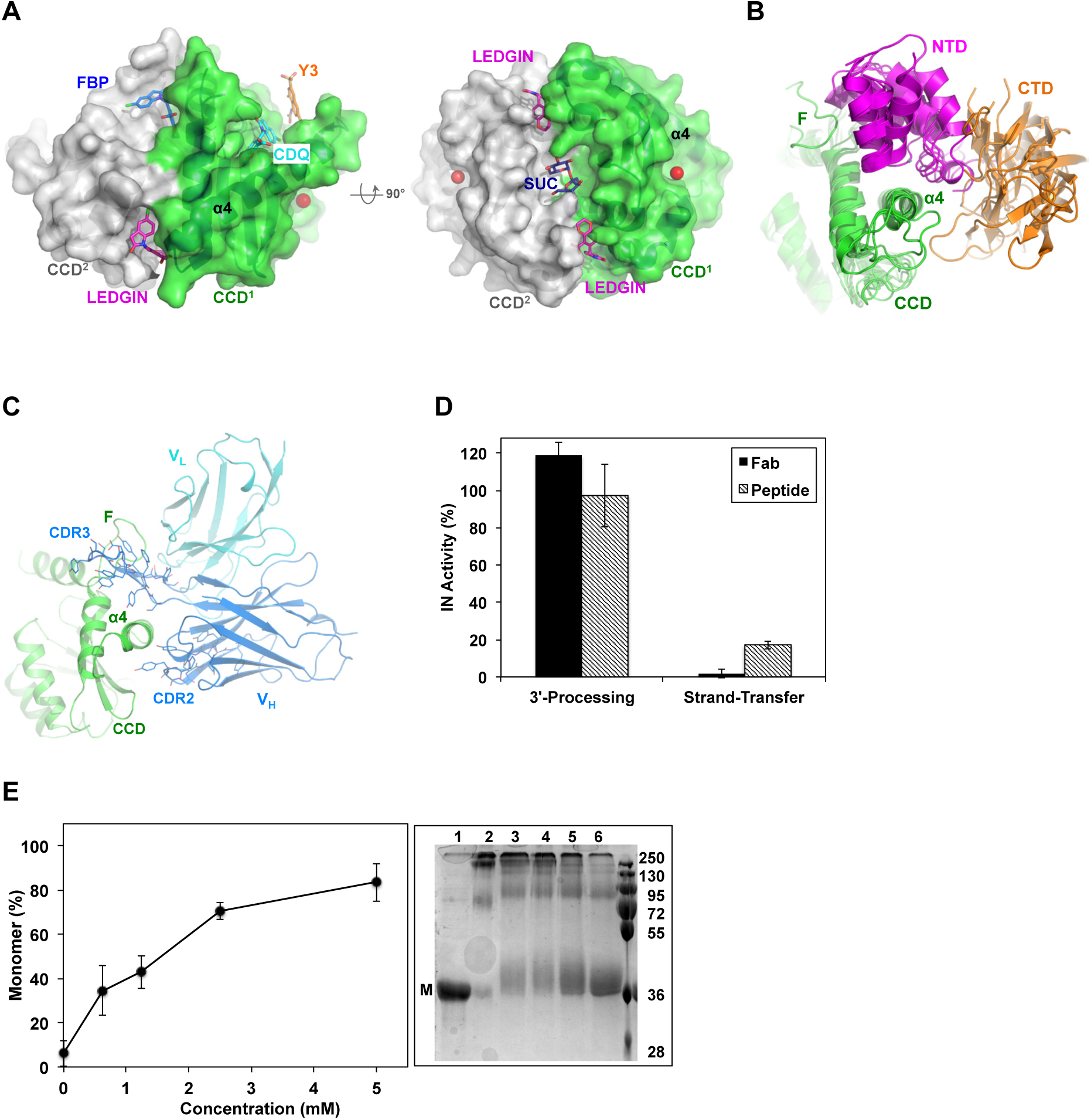
Allosteric sites of HIV-1 IN and the inhibition of IN activity. **(A)** Surface rendering of HIV-1 IN CCD dimer (PDB code: 3NF8) showing the various allosteric sites. LEDGIN (magenta sticks), FBP (blue sticks) and Y3 site bound to CDQ molecule (cyan sticks, PDB code: 3NF8) and Y3 molecule (orange sticks, modeled from PDB code: 1A5V). Sucrose (SUC, blue sticks, PDB code: 3L3V) is also modeled into the structure. Magnesium ion (red sphere) was superimposed from HIV-1 intasome (PDB code: 5U1C) to implicate the catalytic site. **(B)** Conservation of NTD (magenta) and CTD (orange) packing around α4 of CCD (green) at site-1. Structures are of HIV-1, MVV, RSV, MMTV and PFV intasomes (PDB codes: 5U1C, 5M0R, 5EJK, 3JCA, 3OS0, respectively). **(C)** Structure of IN CCD (green) bound to CDR3 and CDR2 of variable heavy (V_H_, blue) and light (V_L_, cyan) chains of Fab specific to site-1 (PDB code: 5EU7). Interacting Fab residues are shown with lines. F: finger loop. **(D)** IN catalytic activity as affected by Fab and the inhibitor peptide. Values are means of three repeats (± SEM). **(E)** Effect of peptide binding on IN multimerization. Values are means (± SEM) of band intensities from three SDS-PAGE gels resolving the various multimeric states of IN (right panel). Representative gel resolving the monomeric (M) form of uncross-linked sample of full length IN (lane 1) or cross-linked in the absence (lane 2) or presence of increasing concentrations of peptide (lanes 3-6). Molecular mass standards (kDa) are shown.

Based on the structure of Fab specific to IN CCD/NTD platform (9), we develop a peptide that, similar to Fab, inhibits IN multimerization and strand-transfer activity. Surprisingly, crystal structure and affinity experiments show that the peptide interferes with the CCD/CTD interfaces of IN. This structure, which features an overlooked SH3-docking platform crucial for IN multimerization, can now provide a template for the screening and development of novel anti-IN allosteric inhibitors.

## RESULTS

### Fab derived peptide inhibits IN multimerization and strand-transfer activity

Within a functional IN tetramer, the NTD and CTD of one dimer wrap around an extended α-helix (α4) of CCD of a second dimer (Figure 1B), an interdomain-docking platform that is functionally imperative (9). Previously, we showed that complementarity-defining regions 3 (CDR3) and 2 (CDR2) of an anti-IN Fab wrap around α4 of IN CCD (residues 153-168) and dock into a cleft rimmed by the CCD finger loop (residues 186-191) (Figure 1C). We derived a peptide (W<u>SYFYDGSYSYYDYE</u>SY) mostly representing the CDR3 sequence (underlined) with the addition of Trp (for UV-Vis absorbance) and Ser-Tyr (representing CDR2 extension in wrapping around α4). Similar to parental Fab, the peptide inhibited IN strand-transfer activity but not 3’-processing (Figure 1D), suggesting potential interference with IN functional multimerization.

To evaluate the effects of the peptide on IN multimerization we used chemical crosslinking and found the peptide to indeed interfere with IN multimerization (Figure 1E). This underscores the allosteric potentials of the site(s) targeted by the peptide, and clarifies that the inhibition of IN multimerization and activity by the parental Fab (9) was specific to CDR3/CDR2 local allosteric interference and not an artifact of the bulkier size of whole Fab.

### The peptide targets two SH3-docking sites at the CCD/CTD interfaces

To characterize the peptide binding site(s), we solved the 2.0 Å crystal structure of HIV-1 IN CCD in complex with the peptide (table-1). Except for the C-terminal pair of residues (Ser-Tyr), well-defined electron densities show binding of the peptide molecule to two sites, which we call site-1 and site-2 (Figure 2A). By examining available structures of various IN functional multimers (4–8) we underscore the conservation of these two CCD/CTD docking platforms (Figure 2B). Site-1 is at the tetrameic interface and is formed by a CTD from a flanking protomer docking at the carboxyl-terminus of CCD α4 of an inner protomer (Figure 2B & 1B). Forming site-2 is the docking of a CTD of another flanking protomer into the amino-terminus of the same α4 (Figure 2B & 2C). The surface area buried upon peptide binding at site-2 is ~ 100 Å^2^ larger than that at site-1 (744 as compared to 650 Å^2^). Peptide binding at site-2 also involves a higher number of interacting residues at the interface (26 as compared to 17 at site-1). Side chains of two residues of the peptide (Trp1 and Tyr3 at site-1 and Ser10 and Tyr14 at site-2) participate in direct hydrogen bonds with the CCD (to side chains of Glu69 and Asp167 at site-1, and to carbonyl oxygen of Val79 and Ala80 at site-2). Additional hydrogen bonds were directly formed with the backbone of the peptide (one in site-1 and three in site-2), and water molecules mediated additional interactions at each site (two in site-1 and one in site-2) (Figure 2D). Interestingly, Ala80 carbonyl oxygen of CCD at site-2, which makes a hydrogen bond with the peptide, also makes a hydrogen bond (2.93 Å) with Lys266 of CTD in the HIV-1 IN functional multimers (discussed below and Figure 4B). It has been shown that the positive charge at this Lys266 position is crucial for viral replication in which a Lys266Ala or Lys266Glu produced replication defective viruses (23).

**Table-1:** Crystallography data collection and refinement statistics

|  | HIV-1 IN-Peptide |
| --- | --- |
| **Data collection** |  |
| Space group | P 41 21 2 |
| Mol/ASU | 1 |
| Cell dimensions |  |
| *a*, *b*, *c* (Å) | 96.52 96.52 48.49 |
| $\alpha$, $\beta$, $\gamma$ (°) | 90 90 90 |
| Resolution (Å) | 68.25-2.01 (2.09-2.01)* |
| *R*<sub>sym</sub> or *R*<sub>merge</sub> | 0.061 (0.885) |
| *I* / *sI* | 17.9 (2.6) |
| Completeness (%) | 99.96 (99.93) |
| Redundancy | 9.7 (10.2) |
| **Refinement** |  |
| Resolution (Å) | 2.01 |
| No. unique reflections | 15680 (1524) |
| *R*<sub>work</sub> / *R*<sub>free</sub> (5% test set) | 0.225 (0.351) / 0.279 (0.364) |
| No. atoms |  |
| Protein | 1176 |
| Water | 51 |
| *B*-factors |  |
| Protein | 47.87 |
| Water | 52.51 |
| R.m.s. deviations |  |
| Bond lengths (Å) | 0.007 |
| Bond angles (°) | 0.84 |
| Ramachandran (%) |  |
| Favored | 95.92 |
| Allowed | 4.08 |
| Outliers | 0.00 |
| PDB code | 6EX9 |

\* Number in parentheses is for highest resolution shell.

**Figure 2.**
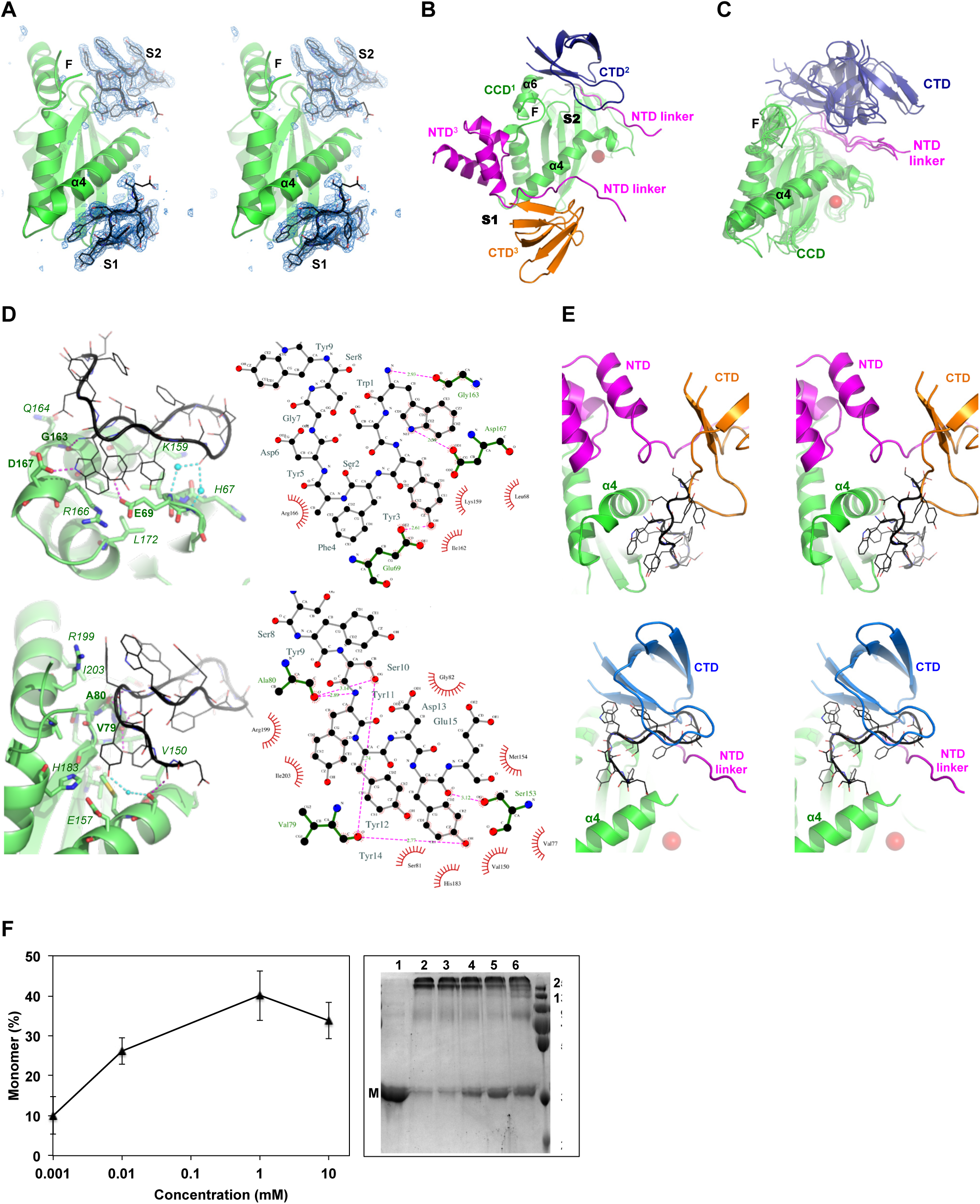
Crystal structure of IN CCD-peptide complex. **(A)** Walleye stereo view of CCD (green) with the peptide and its symmetry mate bound at site-1 (S1) and site-2 (S2), respectively. Blue mesh represents electron density maps (F_O_-F_C_, 3.0σ) calculated after omitting the peptide residues (black). **(B)** NTD (magenta) and CTD (orange) docking at site-1 (S1) and CTD (blue) docking at site-2 (S2) within the same CCD α4 (green). F: finger loop. Red sphere: magnesium ion of catalytic site (PDB code: 5U1C). **(C)** Conservation of CTD (blue) packing against NTD linker (magenta) and α4 of CCD (green) at site-2. Structures are of HIV-1, MVV, RSV, MMTV and PFV (PDB codes: 5U1C, 5M0R, 5EJK, 3JCA, 3OS0, respectively) intasomes. F: finger loop. Red sphere: magnesium ion of catalytic site (PDB code: 5U1C). **(D)** Analysis of peptide (black) interactions with the CCD (green) at site-1 (top panel) and site-2 (bottom). Interacting residues are shown with lines (peptide) or sticks (CCD). Dashed lines indicate direct hydrogen bonds (magenta) or mediated (cyan) by water molecules (cyan spheres). **(E)** Walleye stereo view of the HIV-1 intasome (PDB code: 5U1C) superimposed with the peptide (black) bound to site-1 (top) and site-2 (bottom). Red sphere indicate magnesium ion of catalytic site. **(F)** IN multimerization as affected by the Y3-molecule and is presented by the percentage of monomer formation. Values are means (± SEM) of band intensities from three SDS-PAGE gels resolving the various multimeric states of IN (right panel). Representative gel resolving the monomeric (M) form of uncross-linked sample of full length IN (lane 1) or cross-linked in the absence (lane 2) or presence of increasing concentrations of Y3-molecule (lanes 3-6). Molecular mass standards (kDa) are shown.

Whereas the peptide mainly represented the CDR3sequence and was expected to also represent CDR3 binding pattern (to dock at the CCD/NTD interface between α4 and the finger loop, (Figure 1C)), the peptide surprisingly docked underneath α4 to mostly resemble the binding of CDR2 at site-1 (Figure 2A). Superimposing the CCD-peptide structure onto that of the HIV-1 intasome reveals potential steric interference with the CCD docking platform of CTD but not NTD at site-1 (Figure 2E, top). Similarly, the symmetry related molecule of the peptide bound at site-2 reveals potential interference with the docking of another CTD (Figure 2E, bottom).

To demonstrate the ability of a fragment specific to site-2 to interfere with CCD/CTD docking and consequently inhibit IN multimerization, we evaluated the inhibitory effects of the Y3-molecule, which specifically binds at site-2 (16). Indeed, using chemical crosslinking we show that the Y3-molecule attenuates IN multimerization in a dose-dependent manner (Figure 2F), corroborating on the potentials of a site-2 bound peptide to inhibit IN multimerizaion.

Therefore, peptide binding at either of these sites, or both, would inhibit CTD interactions with CCD, and can rationalize the observed inhibition of IN multimerization (Figure 1E) and strand-transfer activity (Figure 1D). It is worth noting that since both docking sites involve not only the CTD but also the NTD linker segment (residues 50-58) (Figure 2B), peptide binding would also interfere with the vital docking of NTDs during IN multimerization.

Competency of peptide-bound IN for 3’-processing activity indicates that peptide binding did not restrict the catalytic loop flexibility (density map is missing for residues 140-147) nor induced inactivating conformational changes to the enzyme (root-men-square deviation [RMSD] 0.30 Å to apo, PDB code: 1BIS).

### SH3-docking platform-2 offers an ideal spot for allosteric targeting

The higher number of interactions and the larger surface area buried upon peptide binding to site-2 may highlight a binding privilege at this site.

Analyses of crystal structures of IN truncation variants, which have been shown to dimerize (CCD and CCDCTD) or tetramerize (NTDCCD) in solution and crystals, show the clear exposure and accessibility of site-1 (Figure 3A and B). Whereas site-2 is also exposed in the crystal structures of dimeric CCD and tetrameric NTDCCD (Figure 3A), analyses of CCDCTD crystal structures intriguingly show that CTD is constantly docked into site-2, albeit in various configurations distinct to that of the functional intasome (Figure 3B, orange cartoon). If indeed the CTD preoccupies site-2 within the truncated CCDCTD variant, as seen in crystal structures, then we expect that, unlike CCD or NTDCCD variants, the CCDCTD construct would not bind the peptide. Assessing peptide-binding affinities to the various IN constructs show that the presence of NTD (within NTDCCD) only slightly (~ 1.5 folds) interferes with peptide binding (Figure 3C). CTD (within CCDCTD), on the other hand, almost completely abolishes peptide binding to more than 12 folds (Figure 3C). In agreement with the binding data, structural superimposition of the peptide shows that while NTD (of NTDCCD structure) may delicately interfere with peptide binding at site-1 (Figure 3A), CTD (of CCDCTD structures) would provide a more robust interference at site-2 (Figure 3B). It is worth noting that the peptide bound to full-length IN, which is preassembled into tetramers in solution (24), with affinities (Kd 1700 ± 88 nM) corresponding to the concentrations in the monomerization assay (Figure 1E) and ~ 4 folds weaker than NTDCCD (Kd 441 ± 11 nM, Figure 3C). Together, the NTD appears to crucially modulate CTD interactions with the CCD.

**Figure 3.**
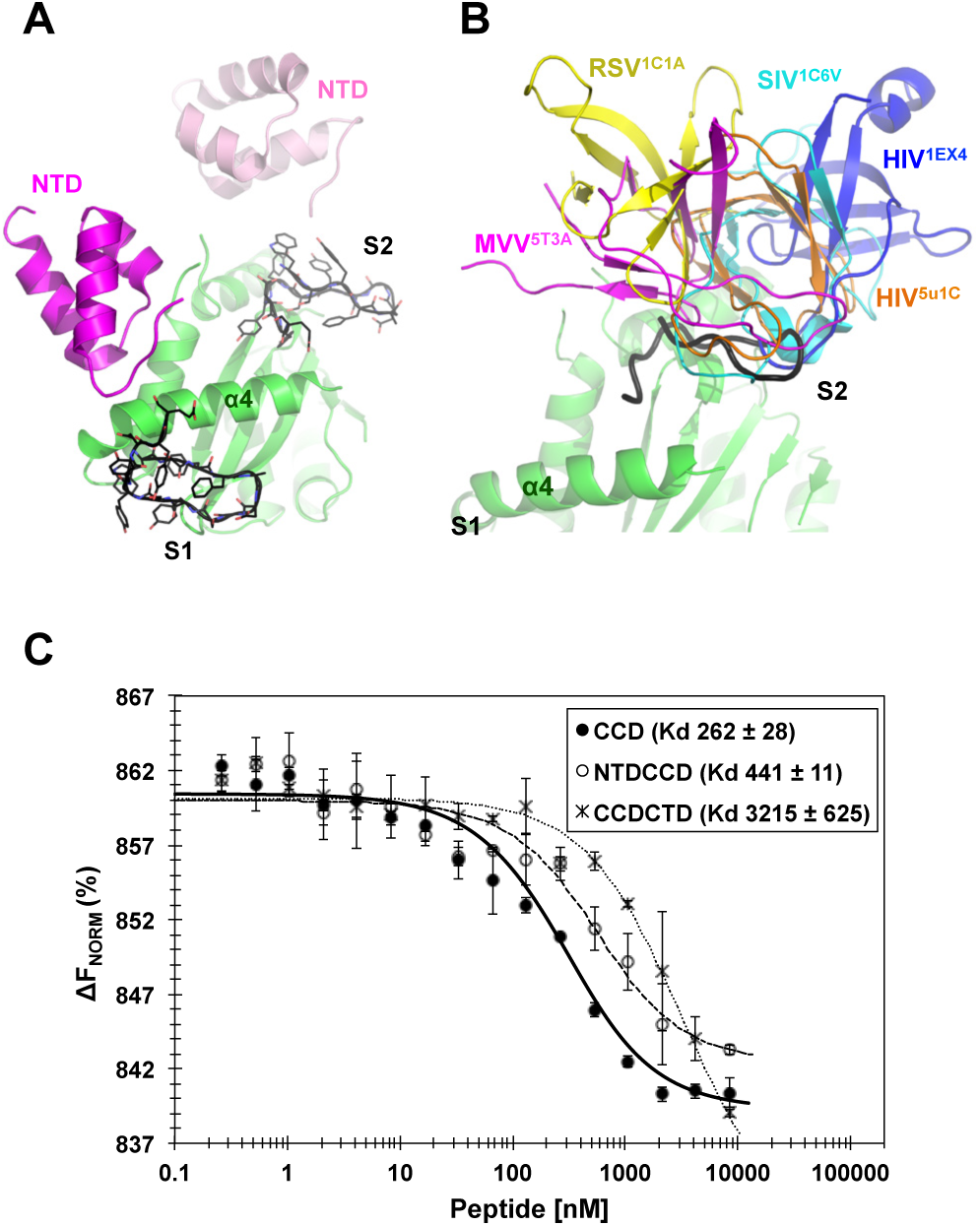
Peptide binding affinities to IN truncation domains. **(A)** Superposition of the peptide (black), bound at site-1 (S1) and site-2 (S2), to NTDCCD structure (PDB code: 1K6Y). **(B)** Superposition of the peptide (black), bound at site-2 (S2), to the various CCDCTD structures of HIV-1 intasome (orange, PDB code: 5U1C) or apo proteins of HIV-1 (blue, PDB code: 1EX4), SIV (cyan, PDB code: 1C6V), RSV (yellow, PDB code: 1C1A), and MVV (magenta, PDB code: 5T3A). CTD of 1EX4 is modeled from a symmetry mate. Site-1 (S1) of all CCDCTD structures is clearly exposed and therefore a peptide was not superposed at S1. **(C)** Peptide binding affinity to HIV-1 IN CCD, NTDCCD and CCDCTD. Values are means of three repeats (±SD). ΔF_Norm_: normalized fluorescence units (% of bound and unbound peptide).

### Implications for the CTD interdomain interactions

The CTD preoccupation of site-2 in the two-domain CCDCTD construct may have significant implications on both the physiologic and the aberrant interactions mediated by the SH3-containing CTD domain. The incompetence of CCDCTD variant for 3’-processing activity (25,26) indicates that the CCD/CTD configuration seen in the crystal structures of this construct (Figure 3B) is aberrant. Conceivably, interdomain interactions, especially those provided by the NTD and missing in CCDCTD, are required for correct CTD interactions and to maintain the catalytic site of the enzyme accessible for catalytic activities.

To gain further insights into the interactions made by NTD and CTD, and their interference with the catalytic activity of CCD, we assessed 3’-processing activity of truncated domain variants of HIV-1 IN. These truncation variants are not active in strand-transfer activity (24–26), a result that we also validated here (not shown). We found that whereas the CCD (residues 56-209) and NTDCCD (residues 1-209) variants of HIV-1 IN are as competent for the 3’-processing activity as the full-length enzyme, the CCDCTD (residues 56-288) is indeed defective (Figure 4A).

**Figure 4.**
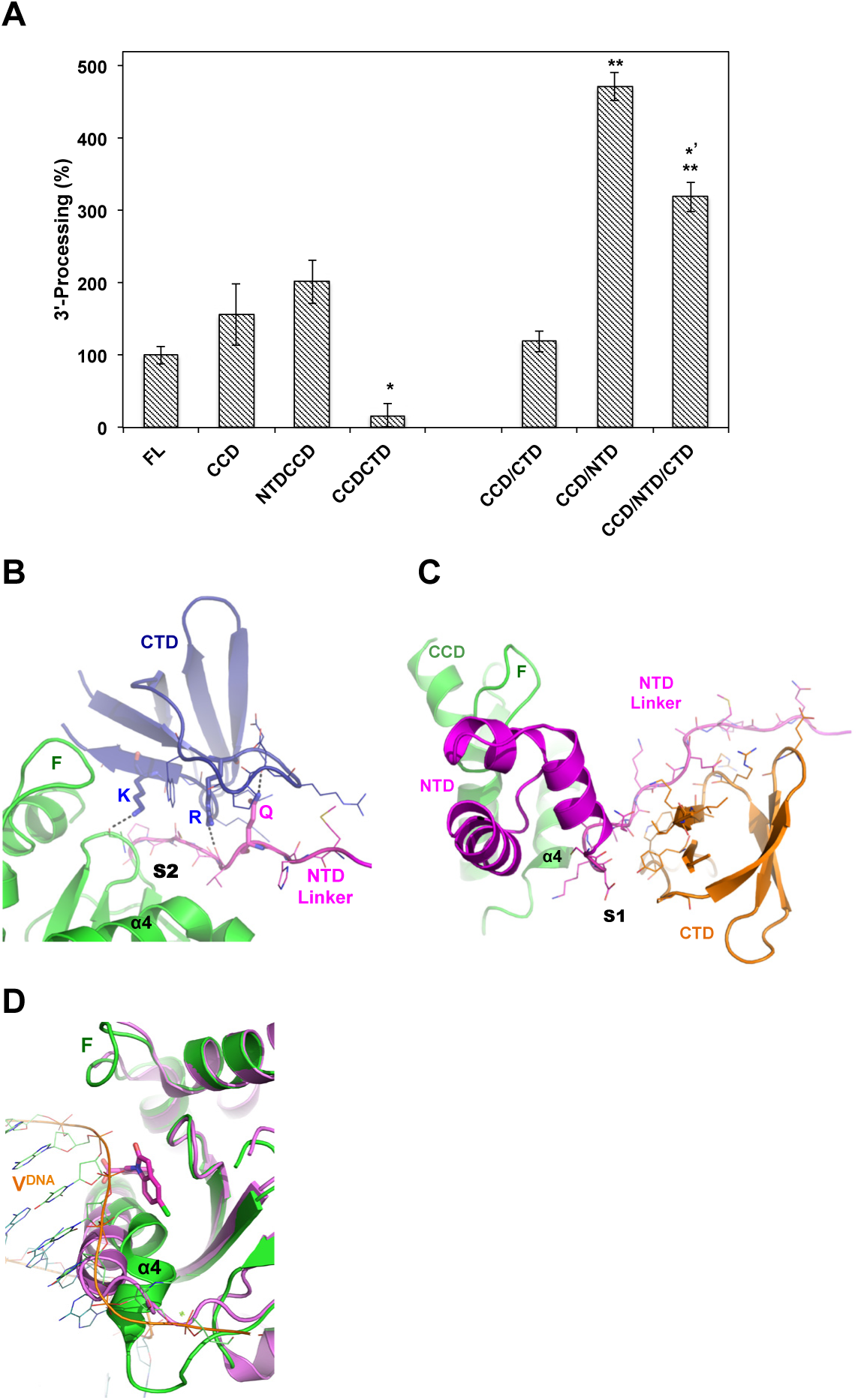
IN activity and multimerization. **(A)** 3’-processing activity (%) of full-length HIV-1 IN (FL) and truncation variants. Whereas NTDCCD indicate two-domain constructs, CCD/NTD indicate that single domain protein CCD and single domain NTD were purified separately and mixed together in the reaction. Values are means of three repeats (±SEM). P-values (one-way ANOVA) are < 0.05 (*) or < 0.01 (**) as calculated in comparison with FL sample, or < 0.05 (*’) as compared to CCD/NTD. **(B)** CTD (blue) interactions with CCD (green) or NTD linker (magenta) at site-2 (S2). Dashed lines indicate hydrogen bonds made by K266 (K) or R228 (R) or CTD and Q53 (Q) of NTD. F: finger loop. Interacting residues are shown with lines. **(C)** NTD (magenta) and CTD (orange) docking at site-1 (S1) of CCD (green). F: finger loop. Interacting residues are shown with lines. **(D)** Structure of CCD (magenta, PDB code 3NF8) in complex with CDQ molecule (magenta sticks) bound at site-2. Superimposed is the CCD (green) of HIV-1 intasome (PDB code: 5U1C). Viral DNA (V^DNA^) of HIV intasome is shown in orange. Binding of CDQ molecule displaces α4 by ~ 5 Å (measured at G149). F: finger loop.

The inhibitory effect of CTD, as part of the CCDCTD, was diminished upon separating the CTD segment (residues 210-288) and supplementing it to the CCD protein in the reaction mixture (Figure 4A, CCD/CTD). Furthermore, the addition of CTD to a reaction containing the separate CCD and NTD (residues 1-55), which by itself (Figure 4A, CCD/NTD) was found more than 3 folds superior to full-length or conjugate NTDCCD, resulted in a significant 33% inhibition (Figure 4A, CCD/NTD/CTD). Therefore, while NTD positively interferes with the CCD in the 3’-processing activity, the CTD appears to have a negative effect. The contribution of the NTD appears necessary to support appropriate CTD interactions with the CCD. Structural analysis shows that the NTD linker segment (residues 50-58) contributes 55% (725 Å^2^) of the total 1310 Å^2^ buried CCD/CTD area at site-2 (Figure 4B). Interestingly, the carbonyl oxygen of Val54 of NTD linker forms a hydrogen bond (2.91 Å) with the side chain of CTD Arg228 (NH2) (Figure 4B), mutating of which to Arg228Ala resulted in defective HIV-1 viruses (23). The side chain of NTD Gln53 (NE2) forms a hydrogen bond (2.50 Å) with the carbonyl oxygen of CTD Asp229 (Figure 4B), and also interacts with the side chain of Lys264 (3.6 Å). Both Lys264 and Lys266, which hydrogen bond to Ala80, have been shown to play an important role in the LEDGINs induced aberrant multimerization (27). Indeed, the fundamental role of CTD in aberrant IN aggregation has previously been emphasized (10,27,28). CTD interactions with the NTD linker (residues 44-53) at site-1, which provides the main docking template for CTD at this site, buries a total of 944 Å^2^ (Figure 4C). Therefore, the NTD linker appears to play a crucial role in modulating the CCD/CTD docking platform and functional IN mutlimerization and is worth exploring in future studies.

## DISCUSSION

Multimerization of HIV-1 IN depends on the delicate compatibility of amino acids lining the interacting interfaces (9), providing an attractive targeting strategy. Interfering with the interacting domains can be exploited not only to promote aberrant multimerization but also to inhibit the assembly of functional multimers, an opportunity offered by the CCD/NTD (HTH-cleft) (9) and the CCD/CTD (SH3-cleft) docking platform reported here. The SH3-cleft (site-2) relates to a previously identified “Y3” pocket observed almost two decades ago in a crystal structure of retroviral IN CCD bound to the small molecule (Y3) (Figure 1A), which efficiently inhibits IN catalytic activity (16). A recent fragment-based screening study has also captured several other small molecules bound near this site (21) (Figure 1A, CDQ), however, the inhibitory profiles where not characterized.

The inability of peptide to bind CCDCTD, which possesses an exposed site-1 but blocked site-2, indicates that site-2, which docks the CTD during multimerization, is a preferred binding site. CTD involvement in the multimerization of IN is well documented (reviewed in (23)), and the relative orientation of the CTD with regard to the CCD has previously been implicated in functional IN assembly (29). Moreover, analysis of available CCD structures bound to small molecules at site-2 reveals a pocket that forms upon binding, which displaces the flexible N-terminus of α4, a displacement that would also interfere with and inhibit the binding of viral DNA and IN catalytic activity (Figure 4D). Therefore, we propose site-2 for future structure based design or screening of small molecules that can interfere with CCD/CTD interactions during IN multimerization, and can promote the formation of a binding pocket that would also inactivate the catalytic site by displacing α4 into an aberrant configuration.

The fact that IN regulates not only viral integration but also the assembly and maturation of virus particles (9,22), accentuates the prominence of this target. Whereas the peptide used in this study may not, *per se*, provide a lead hit for this anti-IN strategy, our work underscores the overlooked SH3-docking platform of HIV-1 IN as a potential therapeutic target for future anti-IN allosteric inhibitors.

## EXPERIMENTAL PROCEDURES

### Preparation of HIV-1 IN Constructs

All HIV-1 IN constructs were derived from the previously described variant (SF1: (SF1; C56S, W131D, F185K and C280S (24)) and subcloned into pET28b plasmid (Novagen) with a cleavable N-terminal His-tag. The various constructs contained the following residues: full-length 1-288, NTD 1-55, CCD 56-209, NTDCCD 1-209, CTD 210-288 and CCDCTD 56-288. To include the NTD linker, the NTD 1-55 used here was made longer than our previous construct (residues 1-51 of HFH chimera (9)). All constructs were confirmed by DNA sequencing.

### Protein Expression and Purification

IN constructs were expressed and purified as previously described (24). Lysis buffer contained 50 mM potassium phosphate (pH 7.5), 5 mM Imidazole, 5 mM β-mercaptoethanol and 1M NaCl for the full-length protein or 0.5 M NaCl for all other truncation variants. For the full-length and CTD containing constructs, the lysis buffer was supplemented with 1 mM CHAPS (3-([3-cholamidopropyl]di-methylammonio)-1-propanesulfonate). Purified proteins were dialyzed into a final solution containing 20 mM HEPES (pH 7.4), 1 mM dithiothreitol, and 1 M NaCl for full-length or 0.5 M NaCl for the truncation variants, and 1 mM CHAPS for CTD containing constructs. His-tag was cleaved using Thrombin (2 units/mg, Novagen). A final size-exclusion chromatography step using Superdex-200 (GE Healthcare Life Sciences) was used for all constructs. Protein stocks (30 μM) were stored (-20 °C) in 50% glycerol.

### In Vitro Fluorescence IN Activity Assays

Fluorescence based IN activity assays were performed as previously described (30). Fab was used at a 1:1 molar ratio and the peptide at an excessive ratio of ~ 20 folds.

### IN 3′-processing activity

DNA substrate was prepared by annealing two fragments: 5′-TACAAAATTCCATAGCAGT-*6FAM* and 5′-ACTGCTATGGAATTTTGTA. IN and truncation constructs were added to the reaction mixture to a final concentration of 4.5 μM. Domain combinations were preincubated at a ratio of 1:1 to a final concentration of 4.5 μM and then added to the reaction mixture containing 25 mM Tris-HCl (pH 7.5), 5 mM dithiothreitol and 50 nM DNA, and incubated for 15 minutes on ice. Next, MnCl_2_ (10 mM) was added and the reactions were incubated at 37 °C for 1hr, and quenched by adding SDS (0.25%). Reaction products were purified using Q beads (GE Healthcare).

### IN single-site integration assay

Donor DNA was prepared by annealing 5′-TACAAAATTCCATAGCA and 5′-ACTGCTATGGAATTTTGTA-*6FAM*, and the acceptor DNA was annealed from 5′-*Biotin*-TATCCGCGATAAGCTTTAATGCGGTAG and 5′-*Biotin*-CTACCGCATTAAAGCTTATCGCGGATA. IN and truncation constructs were added to reaction mixtures to a final concentration of 4.5 μM. Domain combinations were preincubated at a ratio of 1:1 to a final concentration of 4.5 μM, and then added to the reaction mixture containing 25 mM HEPES (pH 7.5), 5 mM dithiothreitol, 0.25 μM donor-DNA, 10 mM MnCl_2_ and incubated for 15 minutes on ice. Next, acceptor-DNA was added (1.5 μM) and reactions were incubated at 37 °C for 1hr and quenched by adding EDTA (10 mM). Reaction products were purified using Streptavidin beads (GE Healthcare).

For the inhibition assays of 3’-processing and strand-transfer activities, Fab (4.5 μM, 1:1 molar ratio) or peptide (70 μM, ~ 15:1 excess molar ratio to IN) were added prior to the addition of substrate DNA.

### Crystallization, Data Collection and Structure Determination

Diffracting-quality crystals of protein-peptide mixture (8.8 mg/ml protein supplemented with 1 mM peptide) were obtained in the JCSG-G7 condition (0.1 M Succinic acid pH 7.0, 15% (w/v) PEG-3350). Cryo-solutions were supplemented with 20% (v/v) glycerol and 0.5 M NaCl.

Diffraction data were collected at beamline ID23-1 of the European Synchrotron Radiation Facility (ESRF), and were processed with iMosflm (31). The structure was solved by Phaser molecular replacement (32) using IN CCD (PDB code: 1BIS (33)) as a search model. Electron densities were fitted using Coot (34) and refined in the CCP4 suite (35) using REFMAC5 (36). Structural superposition and figures were prepared using PyMOL Molecular Graphics System (Schrödinger, LLC). Table-1 summarizes data collection and refinement statistics.

### Structural Analysis

Analyses of interactions were performed using COCOMAPS (37) and LigPlot^+^ (38).

### Lysine Crosslinking

Homobifunctional BS^3^ (Pierce, 2.5 mM) was added to full-length IN (20 μM) that was initially mixed with increasing concentrations of inhibitory peptide or Y3-molecule (4-Acetamido-5-hydroxy-2,7-naphthalene-disulfonic acid, (Sigma, S981680)). Reactions were incubated for 30 minutes at room temperature, and quenched by adding Tris-HCl (pH 7.5) to a final concentration of 10 mM and analyzed on SDS-PAGE.

### Affinity Measurements

IN proteins were labeled with Monolith NT protein labeling kit BLUE-NHS (NanoTemper Technologies) and eluted in microscale thermophoresis (MST) buffer (50 mM Tris-HCl pH 7.0, 300 mM NaCl and 0.05% Tween-20). Labeled IN proteins were incubated with serial dilutions of inhibitor peptide, and MST measurements were carried out using the Monolith NT.115 in standard capillaries with 40% LED and 20% MST power.

## Accession numbers

Atomic coordinates and structure factors were deposited in the Protein Data Bank with the accession number 6EX9.

## Acknowledgments

This work has been supported by iNEXT (grant number 653706, funded by the Horizon 2020 programme of the European Union), the Star Foundation for Applied Research, and benefited from use of the TCSB of Lorry I. Lokey Center for Life Sciences and Engineering and the Russell Berrie Nanotechnology Institute. We thank the European Synchrotron Radiation Facility for the allocation of synchrotron radiation beamtime and the staff for assistance at Beamlines ID23-1.

## Author contributions

M.G. and A.A. designed and performed research, analyzed data and wrote the manuscript.

